# Integrating bacterial and viral genomic information enhances and extends the machine learning predictions of bacteriophage activity against pathogenic *Escherichia coli*

**DOI:** 10.64898/2026.09.23.753777

**Authors:** Antonia Chalka, Ignacio Salinas Valdivieso, Talal Hossain, Christopher Bellas, Alison S. Low, Aidan T. Brown, David L. Gally

**Author notes:** Address correspondence to Aidan Brown, or David Gally. Antonia Chalka and Ignacio Salinas Valdivieso contributed equally to this work. Author order was determined alphabetically by surname.

## Abstract

The use of bacteriophage (phage) to treat bacterial infections is undergoing a resurgence due to the rise of antibiotic resistance. Pathogenic *Escherichia coli,* including extraintestinal pathotypes such as uropathogenic *E. coli* (UPEC), pose a substantial clinical burden with the challenge of multi-drug resistance strains, making it important to progress alternative treatment options such as those based on phage. One major hurdle is selecting effective phage combinations against an infecting strain while accounting for the multiple mechanisms that determine phage susceptibility and bacterial resistance. We previously analysed over 9000 interactions between 31 phage and 314 sequenced *E. coli* and built machine learning models to predict phage activity against ‘unseen’ *E. coli* based on these scores and bacterial genome sequence data. However, this approach did not leverage phage gene content, limiting its ability to generalise across phage. Here, we advance this framework by combining bacterial and phage pangenomes into a single predictive model, allowing gene content across related phage as well as bacteria to be incorporated as machine learning features. This unified approach increased predictive accuracy relative to the single-phage models and produced a single model capable of predicting previously unseen phage–*E. coli* interactions. In leave-phage-out analyses, it also predicted activity for phages whose interaction data were entirely excluded from training. The genomic features most consistently contributing to prediction included bacterial determinants of surface recognition and anti-phage defence, alongside phage-associated features, indicating that this combined model, termed ‘PanPhage’, captures genetic information relevant to multiple stages of the phage–host interaction.

**Importance:** Bacterial infections that once were easily treated with antibiotics are becoming harder to fight as bacteria develop resistance to these drugs. One promising alternative treatment is using bacteriophage (phage), viruses that naturally infect and kill specific bacteria. However, different phages work against different bacterial strains, so ways are needed to quickly match the right phage, or combination of phages, to a patient’s infection.

This study builds a computer model that predicts which phage will inhibit the growth of a given strain of *E. coli*, a bacterium responsible for infections ranging from urinary tract infections to serious bloodstream infections, based on its genetic code. Unlike earlier versions of this model, which only considered bacterial genes, this new approach also includes the phage’s genes. This makes the predictions more accurate and allows activity to be predicted for new phages that were not included during model training, particularly when related phages are represented.

## Introduction

Pathogenic *Escherichia coli* (*E. coli*) is responsible for a substantial share of global bacterial disease, ranging from diarrhoeal illness caused by intestinal pathotypes to urinary tract, bloodstream and other infections caused by extraintestinal pathotypes such as uropathogenic *E. coli* (UPEC) (1, 2). UPEC alone is the leading cause of urinary tract infection (UTI), a condition affecting millions of people annually and increasingly complicated by antimicrobial resistance (AMR)(3). With the rise of AMR, bacteriophage (phage) therapy, the use of viruses to kill bacteria, is undergoing a resurgence as a potential alternative or adjunct to antibiotics (4). Host susceptibility to phage infection is a complex multifactorial relationship, reflecting the continual proliferation of infection, protection and counter protection mechanisms between rapidly evolving phage predators and their bacterial hosts(5). Substantial progress has been made in recent decades in identifying the ways that bacteria can protect themselves against phage infection and the corresponding ways the phage evolve to overcome these defences(5). The challenge for phage therapy is to define phage combinations that act on a particular infecting strain and eradicate it without selecting for a resistant or phage-avoiding bacterial population.

Matching phages to the bacterial strain causing an infection is not straightforward. The first requirement is that the phage can infect the bacterium, which is generally governed by phage-encoded receptor binding proteins (RBPs) and their access to primary and secondary receptors on the bacterial surface (6). These interactions are also impacted by the growth conditions and physiological state of the bacteria, which can substantially alter the efficiency of phage adsorption to their host (7, 8). The number of relevant receptors varies considerably between species: phage infection of *Pseudomonas aeruginosa,* for example, is governed by a relatively limited set of receptors with the great majority of characterised phage using either type IV pili or lipopolysaccharide (LPS) for attachment (9), whereas over 20 distinct surface receptors have been characterised for *E. coli* (6), spanning outer membrane proteins, LPS epitopes, pili and capsule, with considerable strain-to-strain variation further compounded by the polysaccharide capsules that many bacteria produce, depending on the growth conditions (10). Historically, phage selection has relied on laboratory testing of defined phage banks against new bacterial isolates. However, this is specialised and time-consuming work and now sits awkwardly alongside the shift of clinical diagnostics towards genome-based approaches and therefore, a genomic route to phage selection would be desirable.

Any genomics-based approach to phage selection needs to predict the initial ability of a phage to infect a target isolate, and this is reflected in the extensive literature on matching phage RBPs to the bacterial surface targets (11–14). More recently, a number of studies have taken a wider genomics approach to aid phage selection, which may also capture genomic determinants of bacterial resistance acting beyond the initial adsorption step (15–18). The underlying rationale is to capture the genetic diversity of the bacteria and train models on genetic features, such as gene content or short sequence regions (k-mers) alongside phage-isolate interaction data. The resulting models then take unseen test strains and, based on the presence or absence of the specific features, predict whether a given phage can infect them. Available interaction data varies in form, from simple binary plaque formation data through to ‘continuous’ data based on growth curves of isolates with and without phage over time. As with wet-lab phage testing, building appropriate phage-isolate interaction datasets of this kind is time consuming and inevitably constrained by the number of isolates and phage sampled, with substantial datasets likely needed to capture infection and region-specific variation.

A key opportunity is therefore to reduce dependence on extensive interaction datasets by incorporating phage genomic features, allowing predictive information to be shared across genetically related phages rather than requiring each phage to be modelled independently. Our foundation for this approach is our previous work on predictive phage therapy for *E. coli* associated with UTIs, for which over 9000 phage-*E. coli* interactions were measured in an artificial urine medium over 18 hours and scored using an area-under-the-curve calculation based on comparison with growth of non-phage controls (18). This dataset was used to generate individual Random Forest models for each phage, with predictive performance strongly influenced by the balance of susceptible and resistant isolates within the training data. Models for more ‘generalist’ phages, which infected a greater proportion of the *E. coli* collection, generally achieved higher predictive accuracy than those for more specific phages, for which positive interactions were comparatively rare. This was reflected in F1 scores, which provide a combined measure of precision and recall and therefore account for both false-positive and false-negative predictions. While demonstrating that bacterial genomic information could successfully predict phage susceptibility, this approach required sufficient interaction data to train a separate model for each phage, limiting its scalability and its application to newly isolated or poorly characterised phages.

The current work addresses the limitations of individual phage models by incorporating both phage and bacterial genomic features into a single predictive model. By integrating interaction data across phages, the model can share predictive information between genetically related phages, improving the prediction of quantitative phage activity while reducing reliance on extensive interaction data for each individual phage. We show that this approach improves prediction compared with the previous individual phage models and, importantly, can predict activity for genetically related phages excluded from model training, demonstrating the potential to extend predictions to phages with limited or no existing interaction data. Furthermore, identification of the phage and bacterial genomic features contributing most strongly to prediction provides insight into the biological determinants of phage susceptibility, encompassing factors involved in adsorption and infection as well as bacterial resistance mechanisms.

## Results

### Development of the PanPhage model

The primary goal of this study was to extend a previously published phage prediction approach (18), which was solely based on bacterial genomic features, by incorporating phage genomic information into a single combined model, hereafter referred to as the PanPhage model. A simplified overview of the analytical workflow is presented in Fig. 1, with a more detailed version in Supp. Fig. S1.

**Fig. 1:**
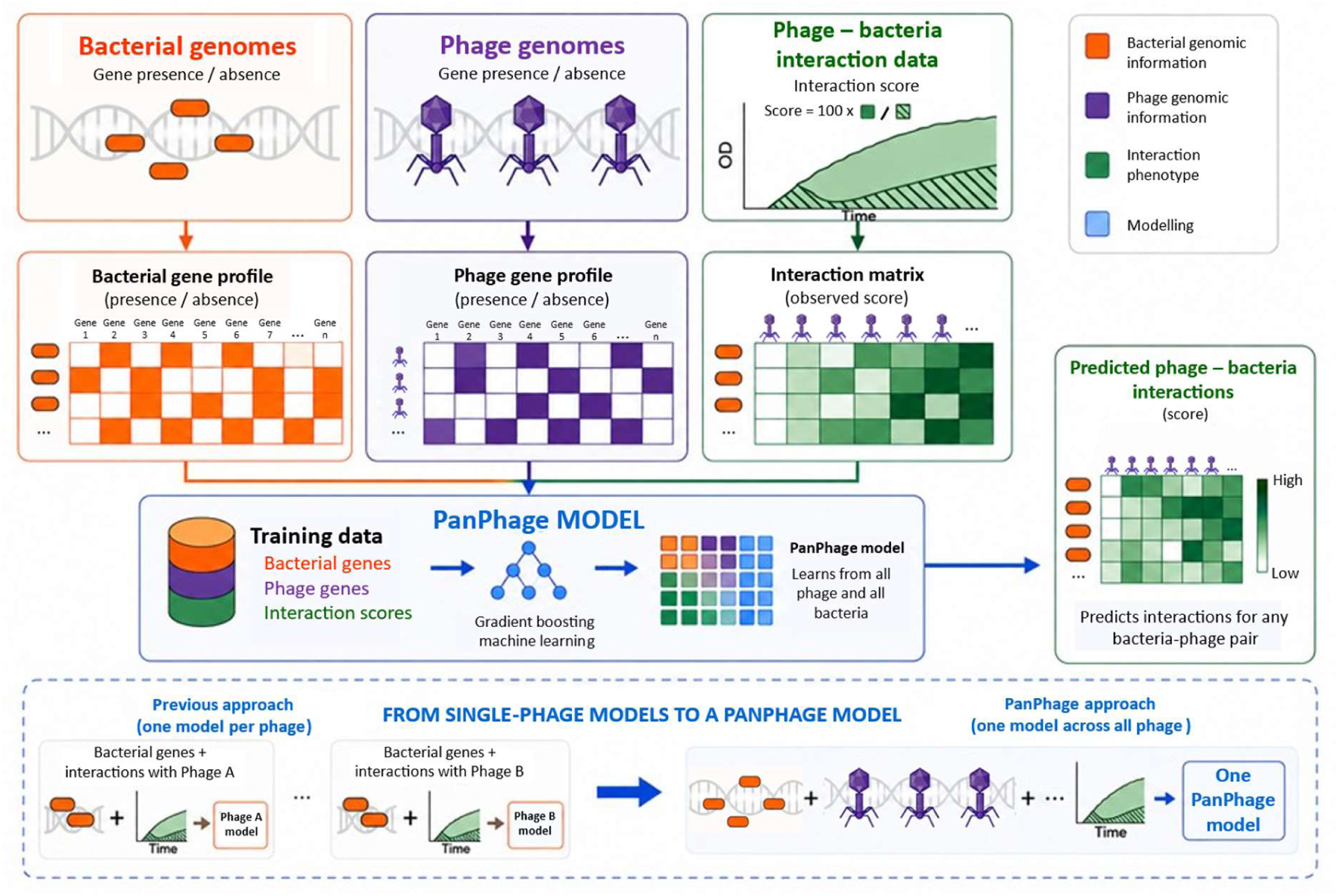
Overview of the PanPhage modelling approach. Bacterial genomic data, phage genomic data and experimentally determined phage–bacteria interaction scores were integrated to train the PanPhage model. Bacterial and phage genomes were represented as binary gene presence/absence profiles, while interaction phenotypes were continuous interaction scores derived from bacterial growth in the presence and absence of phage. These combined features were used to train a gradient boosting model to predict interaction scores for bacteria–phage pairs. In contrast to the single-phage approach, in which an independent model is trained for each phage using bacterial genomic features and interaction data, PanPhage incorporates genomic features from both bacteria and phages into a single model trained across all available phage–bacteria interactions. This enables interaction scores to be predicted for different bacteria–phage combinations.

The interaction dataset used here builds on that described previously (18), in particular using the same set of isolates (phylogenetic tree of *E. coli* strains used in the study is shown in Supp. Fig. S2) and the same raw interaction data. The raw data were reprocessed with two adjustments: outliers resulting from experimental errors were removed, and interaction scores were re-baselined against the mean of the media controls. As described in Methods, interaction scores were calculated from the area under the curve (AUC) for phage-treated wells relative to the no-phage controls, scaled to a 0–100 range, where lower scores indicate effective phage infection and bacterial growth inhibition, and higher scores indicate resistance (heatmap of interaction dataset shown in Supp. Fig. S3 and interaction scores listed in Supp. Table S1). Bacterial genomic features were derived from the previously published genome collection (18), while phage genomic features were newly generated from the 31 phage genome sequences used in this study (Supp. Table S2 and <u>phage genome link</u>). Predicted open reading frames (ORFs) provided the underlying gene content from which binary presence/absence features were derived for machine learning.

We combined bacterial and phage genomic features to train the PanPhage model capable of predicting activity across all bacteria–phage combinations. In parallel, we updated our overall modelling methodology relative to our earlier work (18), including the use of different machine learning frameworks and algorithms. To enable comparison with the earlier single-phage approach, we also retrained the single-phage models (one model per phage) using only bacterial genomic features following the same updated pipeline.

### Overall comparison of single-phage models and the PanPhage model

To evaluate the predictive capacity of the combined genomic approach, we compared the PanPhage model with the retrained single-phage models and null-model baselines that use no genomic features (predicting the mean interaction score from the training interaction data or randomly classifying positive/negative interactions conserving the training data class proportions). Predicted and experimentally measured interaction scores for both the training and unseen test subsets are shown in Fig. 2. Both machine learning modelling approaches showed a clear relationship between predicted and measured scores; however, the single-phage models performed substantially better on the training data than on the test subset (Fig. 2A, C), indicating overfitting to the training data. In contrast, the PanPhage model showed more consistent agreement between predicted and measured scores across the training and test subsets (Fig. 2B, D), with less evidence of overfitting.

**Fig. 2:**
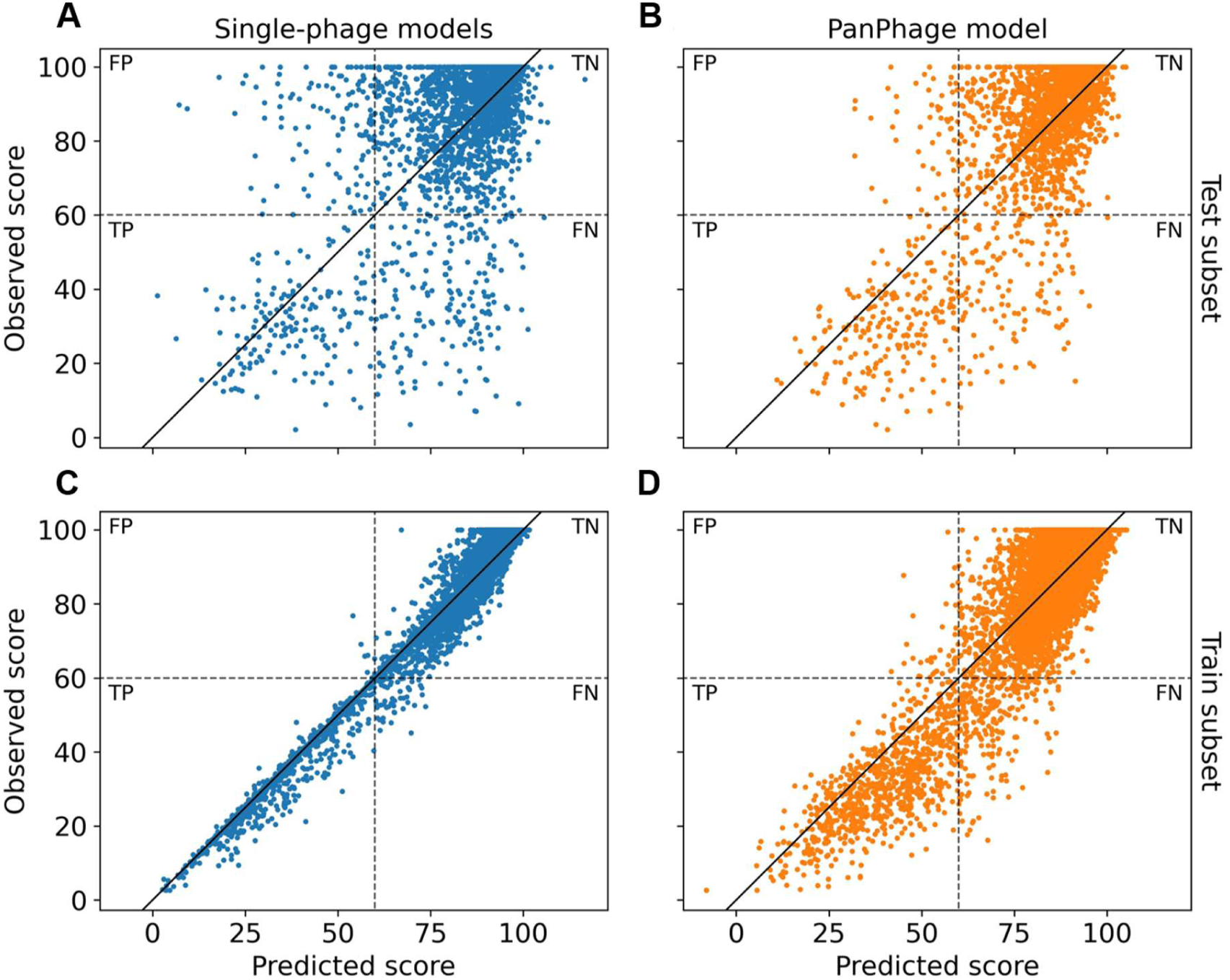
Performance of the single-phage and PanPhage models when predicting interactions in the (A)–(B) test and (C)–(D) train subsets. Each point corresponds to one phage-bacterium interaction predicted by the single-phage models (blue dots) or the PanPhage model (orange dots). Exact predictions lie on the *y* = *x* line (black solid line). The dashed lines delimit the quadrants corresponding to the confusion matrix elements, when a threshold score of 60 is used to define positive (<60) and negative (≥60) interactions: true positives (TP), true negatives (TN), false positives (FP), and false negatives (FN).

The improved performance of the PanPhage approach on unseen interactions was supported by three quantitative performance metrics. For the test subset, PanPhage achieved a root mean square error (RMSE) of approximately 15, compared with 19 for the single-phage models and 22 for the null model. Similarly, PanPhage achieved an F1 score of approximately 0.70 and a Matthews correlation coefficient (φ) of approximately 0.66, compared with 0.52 and 0.45, respectively, for the single-phage models (F1=1 and φ=1 would both correspond to perfect prediction of all binary interaction outcomes). The single-phage models nevertheless substantially outperformed the null-model baseline (F1 = 0.15, φ = −5 × 10^−4^), confirming that bacterial features alone retained meaningful predictive capacity. As expected, the φ coefficient for the null model is approximately zero.

Together, these results demonstrate that incorporating phage genomic features into a combined PanPhage framework improved prediction compared with the independently trained single-phage models.

### Performance of PanPhage across individual phage

We next examined whether the overall improvement achieved by PanPhage was consistent across individual phage and whether performance was influenced by their genomic relatedness within the dataset. PanPhage reduced the RMSE relative to the corresponding single-phage model for all but four of the 31 phage (Fig. 3B), demonstrating that the improvement observed across the complete test dataset was broadly distributed rather than being driven by a small number of phage. The magnitude of improvement varied considerably between phage and was particularly apparent for several phage with close genomic relatives in the dataset. For example, phage within the large related cluster spanning D4 to BO1 and the cluster containing phage A, N, E and P generally all showed substantial reductions in RMSE with PanPhage (Fig. 3B, C). A similar pattern was observed for classification performance, with PanPhage increasing F1 scores for all but four phage for which F1 could be computed (Fig. 3A). For phage ALD1 (* on Fig. 3A), no positive interactions were observed or predicted by any of the models in the test set, reflecting the highly restricted activity of this phage. In this case, the F1 score is undefined, although the models correctly labelled all interactions. Interpretation of absolute RMSE values across individual phage was complicated by substantial differences in their interaction score distributions. As observed previously (18), phage with broader activity generally exhibited greater variation in interaction scores, whereas phage with limited activity produced distributions strongly biased towards high scores. The latter can result in relatively low RMSE values even for models with limited ability to discriminate susceptible from resistant isolates, simply because most observed and predicted scores occupy a narrow range. Consequently, changes in RMSE between the matched single-phage and PanPhage models provide a more informative measure of the effect of incorporating phage genomic information than comparison of absolute RMSE values between different phage. This dependence of RMSE on score distribution may also explain why genomic relatedness was not clearly associated with absolute RMSE, whereas classification performance measured by F1 showed a stronger relationship with phage genomic distance (Supp. Fig. S4).

**Fig. 3:**
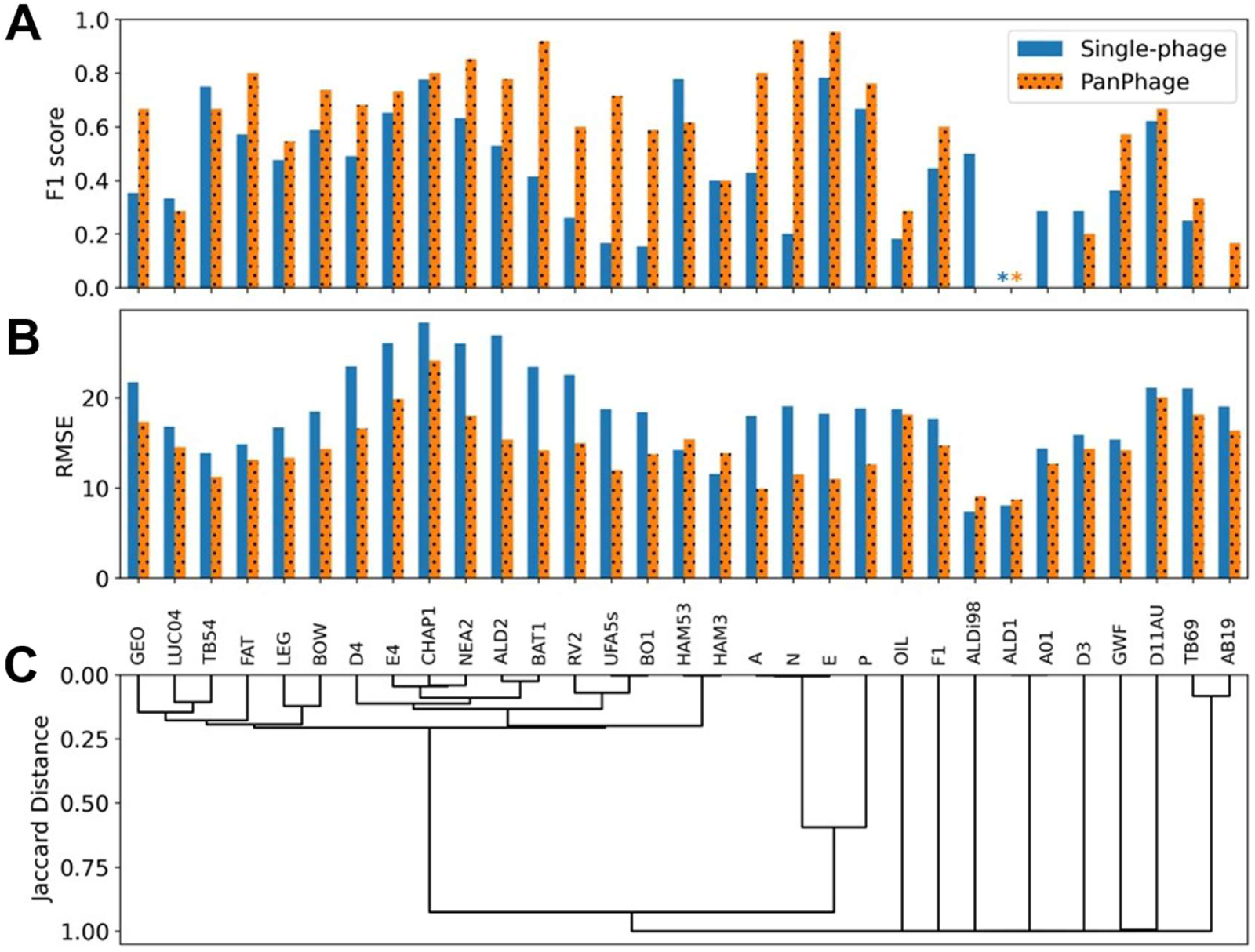
Model performance. measured using **(A)** F1 score and **(B)** Root mean square error for the single phage models (blue bars) and the PanPhage model predicting each phage (orange dotted bars). Asterisks (*) indicate where the respective measure is undefined. **(C)** Dendrogram of the phages in the dataset built using the Jaccard distances in the pangenome matrix. The height of each branch represents the average Jaccard distance between the two clusters above.

### Prediction of interactions for phage excluded from model training

A key potential advantage of incorporating phage genomic features in PanPhage is the ability to transfer information between related phage, potentially allowing predictions to be made for a phage for which no interaction data are available. To directly test this, we performed a leave-phage-out analysis in which all interaction data for each phage were excluded from model training in turn. The resulting model was then used to predict the interaction scores of the excluded phage across the *E. coli* collection, such that prediction relied entirely on information learned from the remaining phage.

PanPhage retained predictive capacity for a substantial proportion of excluded phage, although performance varied markedly across the phage collection (Fig. 4A). This variation was associated with the genomic relatedness of the excluded phage to all the remaining phage. Phage belonging to more closely related clusters, when excluded from training, generally achieved higher F1 scores than did genetically distinct phage with few or no close relatives in the dataset (Fig. 4A, 4C). This relationship was supported by a significant negative correlation between F1 score and the genomic distance of the excluded phage from the remaining phage collection (Pearson’s *r* = −0.59, permutation *p*-value = 0.0005; Fig. 4D). Thus, phage positioned closer in genomic feature space to those represented during model training were more accurately classified when their own interaction data were entirely withheld. In contrast, RMSE showed no clear relationship with phage genomic distance (Fig. 4B, 4C). As observed for the single-phage comparisons above, RMSE is influenced by the underlying distribution of interaction scores for each phage. In particular, phage with restricted activity have scores concentrated towards the upper end of the scale, allowing relatively low RMSE values to be achieved even when the model has limited ability to identify susceptible isolates. F1 therefore provides a more informative measure of whether genomic relatedness supports prediction of biologically relevant phage–bacterium interactions in this analysis.

**Fig. 4:**
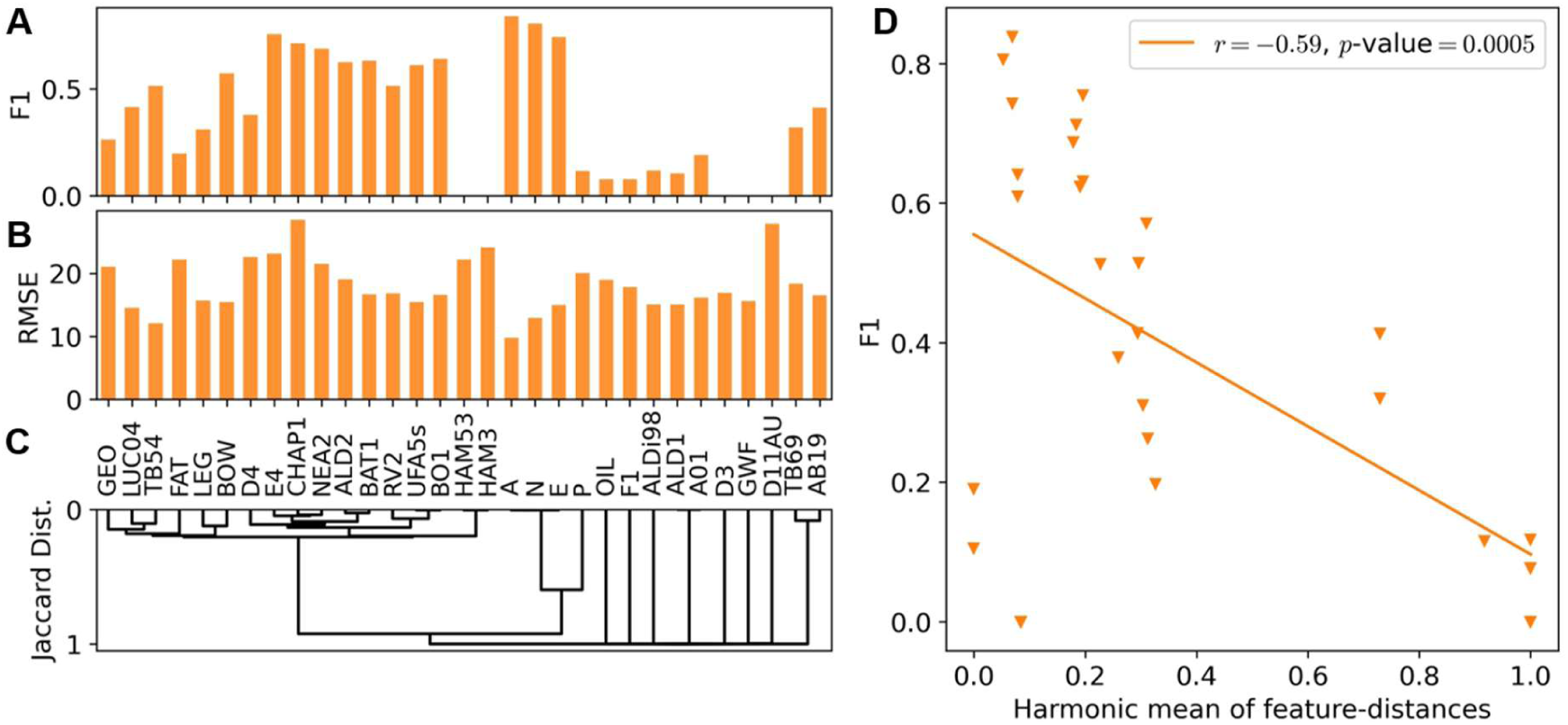
Performance of the Leave Phage Out models predicting the unseen phage, measured using **(A)** F1 score and **(B)** Root mean square error. **(C)** Dendrogram of the phage in the dataset built using the Jaccard distances in the pangenome matrix, repeated from Fig. 3C for convenience. **(D)** Plot of model performance (F1 score) against the harmonic mean of feature-distances of each phage to the rest of the set (using the Jaccard distance in the pangenome matrix).

Together, these results demonstrate that PanPhage can exploit genomic information shared between related phages to predict interactions for phages whose phenotypic interaction data were entirely absent from model training, with predictive performance dependent on the genomic representation of related phages within the training dataset.

### Contribution of bacterial and phage genomic features to model predictions

Having established that the combined PanPhage framework improved predictive performance, we next examined the genomic features contributing to the PanPhage model. To assess the consistency of feature selection, the feature-selection procedure was repeated 128 times using different random seeds (see Materials and Methods for further details), and bacterial and phage features were ranked according to the proportion of these 128 models in which they were retained (Fig. 5).

**Fig. 5:**
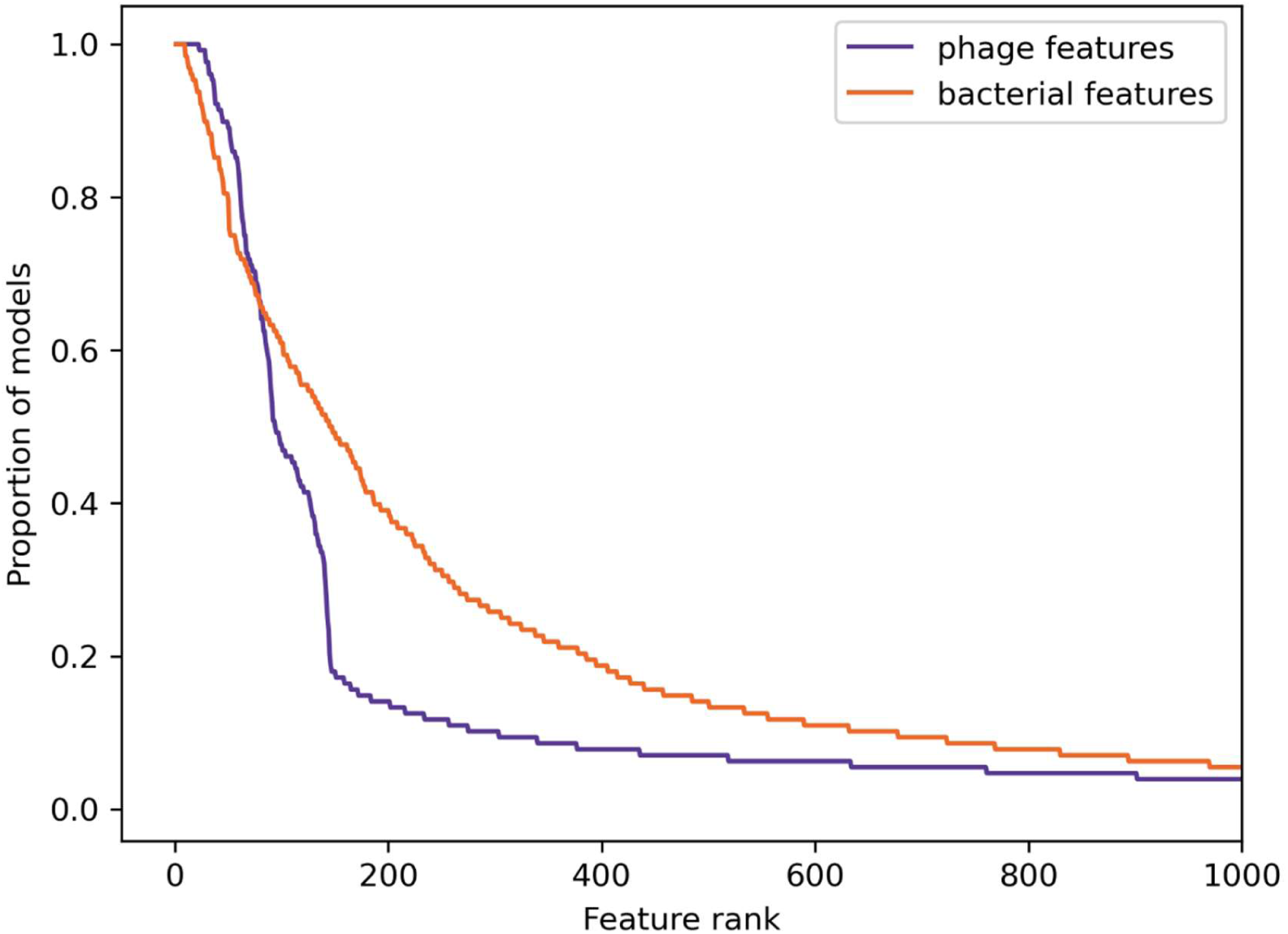
Features ranked by percentage of models that selected them. The ranks of the indices are on the horizontal axis, and the height of the curve represents the proportion of models that select each feature. The purple line corresponds to phage features and the orange line to bacterial features.

Features from both bacterial and phage genomes were reproducibly selected, indicating that the fitted model used information from both feature sets. Phage features showed a steeper selection-frequency profile than bacterial features, with a relatively small subset selected consistently across repeated model runs (Fig. 5). However, the initial phage feature set was substantially smaller than the bacterial feature set (1,779 versus 16,886 features), and therefore direct comparison of the selection-frequency distributions between the two feature classes should be interpreted cautiously. We therefore examined the most frequently selected features in greater detail to determine whether they were associated with known biological determinants of phage–host interactions.

### Frequently selected genomic features and biological interpretation

We next examined which specific genomic features underpinned the model’s predictions, and whether these could offer biological insight into the mechanisms of phage infection and resistance. Recursive feature elimination substantially reduced the number of gene clusters available to the model, indicating that a relatively small subset of genomic features contained much of the information relevant to prediction (Supp. Table S3). We therefore examined features that were reproducibly retained across repeated feature-selection runs to determine whether they could be associated with known mechanisms of phage-host interactions.

Among the bacterial features, 147 were selected in >50% of the 128 feature-selection runs. Approximately 10% of these were associated with capsule biosynthesis, lipopolysaccharide (LPS) or known/potential phage receptors, while a further 8% were associated with anti-phage defence systems. Analysis of gene synteny across the bacterial pangenome showed these features were often clustered across hypervariable regions or on mobile elements.

Capsule-associated features were particularly prominent. Nine frequently selected features occurred across three capsule biosynthesis loci, including five of the six genes within variable region 2 of the K1 capsule locus (Fig. 6A). Features within the corresponding variable regions of the K14 and K7 capsule loci were also selected, providing genomic information capable of distinguishing these capsule types. This is biologically consistent with the established importance of capsular polysaccharides as phage receptors and determinants of host range (19). Features associated with O-antigen biosynthesis were similarly identified in 24 O75 isolates, while a TonB-associated gene was also frequently selected, further supporting the contribution of bacterial surface structures and receptors to model predictions.

**Fig. 6.**
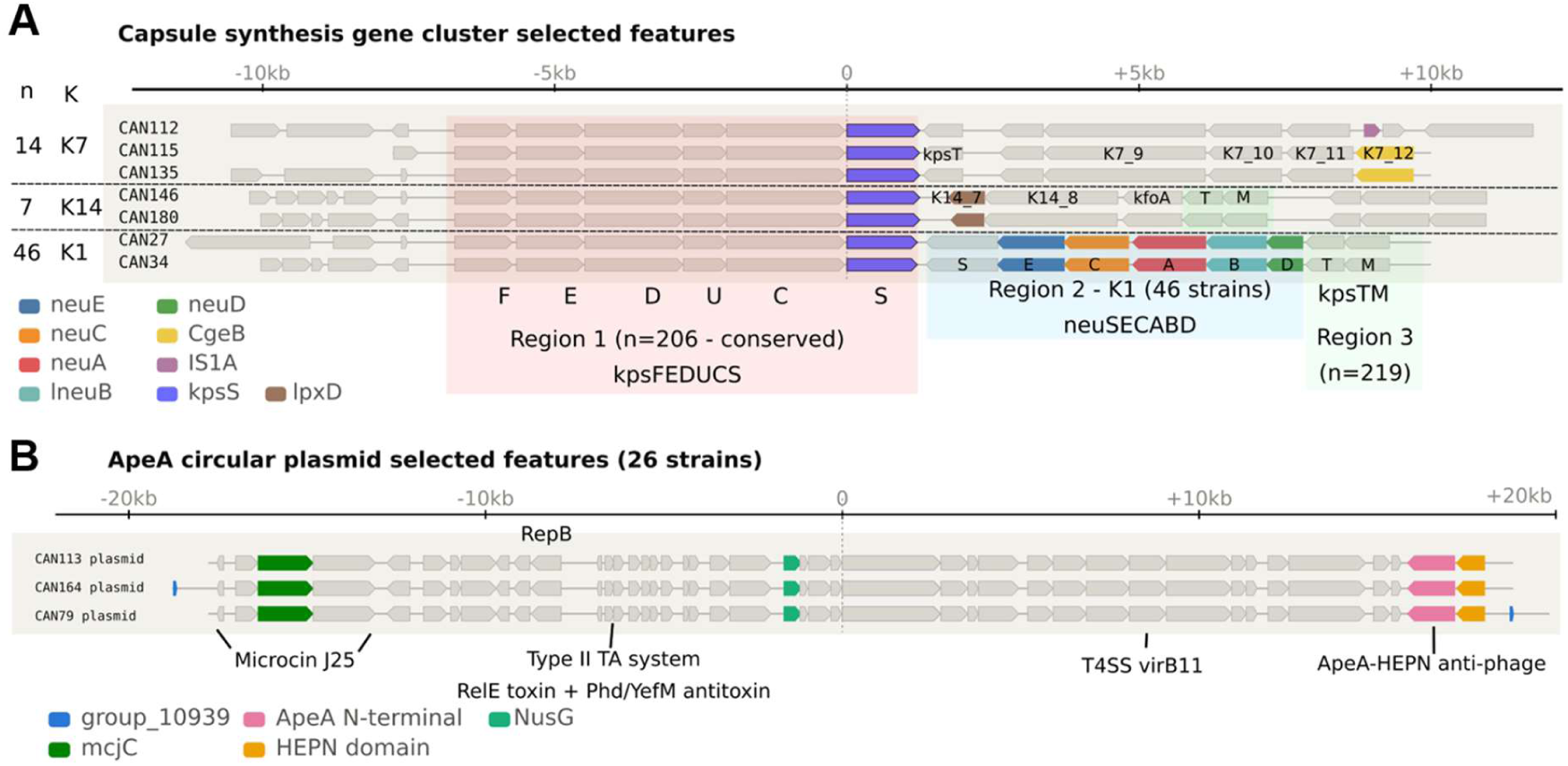
(A) Gene synteny of capsule biosynthesis genes selected by recursive feature elimination. The capsule biosynthesis gene cluster across 67 selected strains was aligned by the conserved region *kpsS*, allowing the selected features (shaded genes) across the hypervariable region 2 to be visualised. n = number of strains the configuration applies to, K = the K-type of the strain, determined by running each contig through kTYPr(20). **(B) Defence plasmid containing ApeA in 26 strains**. The five frequently selected features are highlighted. The circular plasmid is viewed relative to *ptlC* for easy visualisation.

Importantly, frequently selected bacterial features were not restricted to genes associated with phage adsorption. Two restriction-modification systems were represented among the selected features, together with several putative anti-phage defence genes. A mobile element present in 17 isolates was identified through three frequently selected features, including a BstA-family anti-phage gene associated with abortive infection (21). Similarly, four selected features mapped to a 30 kb plasmid present in 26 isolates, including an ApeA HEPN-domain anti-phage gene (22) (Fig. 6B).

Other frequently selected bacterial features were dominated by genes associated with mobile genetic elements, including prophage-associated genes and transposases. Nine features (∼6% of the frequently selected set) were associated with fimbriae, with several corresponding to fimbrial regulatory genes. Although their direct contribution to phage susceptibility is unclear, these features may reflect either additional host-surface determinants or linkage to genomic backgrounds associated with phage susceptibility.

Of the 1,779 phage genomic features, 94 were selected in >50% of feature selection runs. Homing/HNH endonucleases were particularly enriched among these frequently selected features, accounting for 16 of the 94 frequently selected phage features and representing a 5.9-fold enrichment relative to their frequency in the complete phage feature set (Fisher’s exact test *p*-value < 0.001). One homing endonuclease has previously been found to promote phage infection by targeting anti-phage defences; however, whether the enrichment observed here reflects a direct role in infection outcome remains unknown.

Together, these findings indicate that the genomic information used by PanPhage includes bacterial and phage features associated with initial host recognition, phage infection (post entry) and bacterial defence.

## Discussion

This study built on our previous single-phage models for predicting phage activity against pathogenic *E. coli* associated with urinary tract infections (18). The PanPhage model combined bacterial and phage genomic features and outperformed the retrained single-phage models across all metrics assessed (RMSE, 15 versus 19.2; F1, 0.70 versus 0.52; and φ, 0.66 versus 0.45). Improvements were observed across most of the phage collection rather than being driven by a small number of phage. In the leave-phage-out analysis, PanPhage also retained predictive capacity for phage excluded from training, with F1 decreasing as the genomic distance between the excluded phage and those represented in the training data increased (Pearson’s r = −0.59, p = 0.0005).

Representing bacterial and phage gene content within a combined model allowed information to be shared across genetically related phage rather than requiring each phage to be modelled independently from a smaller number of interactions. This is most clearly illustrated by the leave-phage-out analysis, in which phage with close genomic relatives in the training dataset were predicted more accurately than genomically isolated phage. However, PanPhage also differs from the previous single-phage approach in its pooled training dataset, modelling algorithm and feature-selection procedure. The observed improvement therefore reflects the combined PanPhage framework and cannot be attributed solely to the inclusion of phage genomic features. Nevertheless, the leave-phage-out results demonstrate that phage genomic relatedness provides information that can support prediction when interaction data for a particular phage are unavailable.

Modelling phenotypes from bacterial and phage genomes, where the number of potential features greatly exceeds the number of measured interactions, carries a substantial risk of overfitting (25). Limited diversity within the phage and bacterial collections may also result in the selection of features that track genomic relatedness rather than directly determining infection outcome. As previously observed in machine-learning models for source attribution in *E. coli* and *Salmonella Typhimurium* (26, 27), such features may still have predictive value, but they should not be interpreted as causal without experimental validation. Among the 147 bacterial features consistently selected by PanPhage were genes associated with capsule and O-antigen biosynthesis, TonB and known or putative anti-phage defence systems, including BstA and ApeA. These functions are consistent with established determinants acting at different stages of phage infection. However, other selected features, including those associated with fimbriae and mobile genetic elements, may represent linked genomic backgrounds rather than direct determinants of susceptibility.

Homing endonucleases were enriched among the consistently selected phage features, accounting for 16 of 94 features and representing a 5.9-fold enrichment relative to their frequency in the complete feature set (Fisher’s exact test, p < 0.001). Although homing endonucleases can contribute to phage–phage competition during co-infection (24), their relationship with host-range determination is not established. Their enrichment should therefore be regarded as a hypothesis-generating observation that may reflect linkage to other variable regions of the phage genome. Experimental work would be required to determine whether these genes have a direct role in infection outcome.

PanPhage was trained using continuous interaction scores, while classification performance was assessed after applying an activity threshold of 60. This threshold was selected in our previous study asF scores below 60 could be distinguished with high confidence from variation between experimental repeats (18). Nevertheless, the strength of relationships involving F1 may depend on the chosen threshold. The negative association between F1 and phage genomic distance should therefore be interpreted as evidence that closely related phage tend to share infection phenotypes under this classification criterion, rather than as a threshold-independent property of the dataset. By contrast, the weaker relationship between RMSE and genomic distance indicates that prediction of the continuous interaction score is influenced by factors beyond overall phage relatedness, including the distribution and variability of scores for individual phage. Continuous scores retain differences in measured growth inhibition that are lost after binary classification, but the present analysis does not establish whether this additional resolution improves therapeutic phage selection.

Incorporating phage genomic sequence data into the model provided value beyond improved prediction accuracy alone. The pangenome-based clustering of phage revealed clear relationships between genomic relatedness and shared infection phenotypes which will be useful for guiding the selection of phage for inclusion in future interaction datasets or therapeutic cocktails. This approach is complementary to other recent phylogeny-independent strain-level prediction frameworks (28) and, together with recent advances in generative genome design (29), highlights a growing convergence between predictive and generative approaches to phage engineering: features identified as important for infection outcome in models such as PanPhage could, in principle, inform the design or refinement of synthetic phage genomes optimised for activity against specific bacterial targets. Conversely, predictions for phage that are genomically distant from the training collection should be treated with greater caution and prioritised for experimental validation.

This study has several limitations. The phage collection remains modest in size and the bacterial collection is focused predominantly on UPEC and related *E. coli*. In addition, the principal train–test division was performed at the interaction level. Consequently, the main evaluation measures prediction of held-out phage–bacterium combinations but does not necessarily represent prediction for entirely unseen bacterial isolates. The selected genomic features may also include phylogenetic markers, and their biological roles require experimental validation. Despite these limitations, PanPhage demonstrates that combining bacterial and phage pangenome information improves prediction across a densely sampled interaction dataset and enables partial generalisation to phage excluded from training. The framework could therefore be used to prioritise candidate phage for experimental testing while identifying areas of phage genomic diversity for which additional interaction data are needed.

## Materials and Methods

### Bacterial and phage collections

The collection, described previously (18), comprised 314 clinical *Escherichia coli* isolates: 203 from canine urinary tract infections collected at the Royal (Dick) School of Veterinary Studies Hospital for Small Animals between 2017 and 2019, and 111 human isolates collected through the Royal Infirmary of Edinburgh in 2019 (81 urinary and 30 bloodstream isolates). These were designated CAN, HU and HB isolates, respectively.

The 31 phages (Supp. Table S2) were isolated from wastewater supplied by the Scottish Environment Protection Agency or provided by the Phage Technology Centre GmbH (Bönen, Germany). Phage isolation, propagation, titration and storage, bacterial culture conditions and preparation of artificial urine were described previously (18).

### Interaction assays and scores

The previously generated dataset comprised growth measurements for 31 phages tested against 314 *E. coli* isolates in triplicate, with corresponding no-phage controls (18). For the present study, following visual inspection, some technical repeats were excluded from the score calculations because one or more replicate growth curves showed an abnormally high initial signal, a large spike during the initial measurements, prolonged absence of growth in the corresponding no-phage control, or excessive signal noise. The mean signal from negative-control wells was subtracted from each curve.

Area under the curve (AUC) was calculated by trapezoidal integration using SciPy (30). For each phage–isolate combination, the median AUC of the three phage-treated wells was divided by the median AUC of the corresponding no-phage controls and multiplied by 100. Scores were constrained to 0–100, where 0 indicated complete growth inhibition and 100 indicated growth equivalent to the no-phage control.

### Bacterial genome analysis

Whole-genome sequencing and assembly were performed previously (18). Briefly, genomic DNA was sequenced by MicrobesNG using paired-end Illumina sequencing and assembled using the EnteroBase Tool Kit (EToKi). Assemblies containing more than 600 contigs were excluded, leaving 301 *E. coli* genomes; interactions involving the 13 excluded isolates were omitted from model development. EToKi was also used to construct a maximum-likelihood phylogeny from core-genome SNPs.

The 301 genome assemblies were reannotated using Prokka v1.14.5 (31) and the bacterial pangenome generated using Panaroo v1.5.1 (32). Genes were encoded as a binary presence/absence matrix for use as bacterial genomic features.

### Phage genome analysis

Phage DNA was sequenced using 150-bp paired-end Illumina reads. Reads were quality-filtered and adapter-trimmed using Trimmomatic v0.40 (33), then assembled with SPAdes v4.0.0 (34), using k-mer sizes of 27, 47, 67, 87, 107 and 127. Contigs longer than 10 kb were retained. Genome completeness was assessed using CheckV v1.1.1 (35), and assemblies were reoriented using DNAapler v0.8.1 (36).

Protein-coding sequences were predicted using Pyrodigal-gv v0.3.2 (37). Genomes were annotated using Pharokka v1.7.3 (38), followed by structure-informed annotation with Phold v1.2.0 (39). Taxonomic classification was performed using vConTACT3 v3.2.4 and its reference database (40,41). A phage pangenome was generated from the annotated genomes using Panaroo v1.5.1 (32), and genes were encoded as a binary presence/absence matrix.

### Machine Learning models

Gradient Boosting Regressor models were implemented in scikit-learn (41). Single-phage models were trained separately for each phage using bacterial genomic features. PanPhage was trained across all interactions using both bacterial and phage features. Equivalent feature-selection and optimisation procedures and identical train–test assignments were used where applicable.

### Training and test data

Interactions were divided 3:1 into training and test sets at the interaction level. Splitting was stratified by phage and binary activity, with scores below 60 classified as positive interactions and scores of 60 or above as negative interactions (18). This maintained the relative frequency of both classes for each phage and ensured identical test interactions for comparison of PanPhage and the corresponding single-phage models.

### Feature selection

Feature selection was performed using the training data to reduce dimensionality and limit the inclusion of uninformative genomic features that could contribute to overfitting (28). The initial feature sets contained 16,886 bacterial genes and 1,779 phage genes. PanPhage used both feature sets, whereas single-phage models used only bacterial genes. An initial Recursive Feature Elimination (RFE) process was performed, removing the least important features iteratively, based on impurity-derived feature importance, until 2,000 remained (see SI for details). Recursive feature elimination with tenfold cross-validation (RFECV) was applied to these features, removing 200 features per iteration to a minimum of 45 (see SI for details). The number of features giving the best mean cross-validation performance was selected, and RFE was repeated on the complete training set to obtain the final feature set. RFE and RFECV were implemented in scikit-learn (41).

### Hyperparameter optimisation and evaluation

Gradient-boosting hyperparameters were optimised by randomised parameter search with tenfold cross-validation using the training data. The search included the number and maximum depth of trees, learning rate, subsampling fraction, minimum samples required to split an internal node and minimum samples per leaf. The combination giving the lowest cross-validated mean squared error was selected. Search grid, number of sampled combinations and selected parameters are available in the in the SI. The optimised models were evaluated against the held-out test set, which was not used for feature selection or optimisation.

### Performance metrics and null models

Regression performance was assessed using root mean squared error: *RMSE^2^=1/n x ∑_i ∈ Test set_ (ŷ_i_-y_i_)^2^*, where *ŷ_i_-y_i_* are the predicted and observed scores, respectively, and *n* is the number of test interactions. For classification, predicted and observed scores were converted to positive (<60) or negative (≥60) interactions (18). Performance was assessed using F1 score and Matthews correlation coefficient (φ).

The regression null model assigned every test interaction the mean score of the training data; its RMSE was calculated on the test observations. For classification, null predictions were sampled from a Bernoulli distribution with probability equal to the proportion of positive interactions in the training set.

### Association between performance and phage genomic distance

Pairwise phage genomic distances were calculated from the binary gene presence/absence matrix using Jaccard distance:

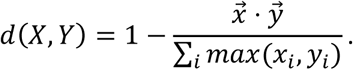

where (X) and (Y) are the sets of genes present in two phages. Each phage’s distance from the remainder of the collection was summarised as the harmonic mean of its pairwise distances.

Pearson correlations were calculated between mean genomic distance and the per-phage RMSE, F1 and φ of the single-phage and PanPhage models. Statistical significance was assessed using 100,000 two-sided permutations (42).

### Feature-selection frequency

To identify features selected consistently despite variation in train–test assignment and model fitting, data splitting and feature selection were repeated 128 times with different random seeds. Selection frequency was defined as the proportion of runs in which each feature was retained.

### Leave-phage-out analysis

Generalisability to phages without training interactions was assessed using leave-one-group-out analysis. In each iteration, all interactions involving one phage were withheld for testing, and the model was trained on interactions involving the remaining phages. Performance for the excluded phage was assessed using RMSE, F1 and φ. Associations between performance and the excluded phage’s genomic distance from the training collection were tested as described above.

### Pangenome synteny plots

Gene-synteny plots were generated using Pansynteny (https://github.com/chrisbellas/pansynteny), with contigs aligned on *kpsS* or *ptlC*, to examine frequently selected features in their genomic context.

## Data and Code Availability statement

### Data

Bacterial genomic information had previously (18) been deposited in Enterobase (https://enterobase.warwick.ac.uk/species/ecoli/search_strains?query=workspace:97040) Phage genomic information has been deposited on figshare (https://figshare.com/articles/dataset/Phage_genomes/33390229). All study data are included in the manuscript or supplementary figures / tables.

### Code

All code used to generate the machine learning models can be found in a github repository – https://github.com/Antonia-Chalka/PanPhage_Models_Reference

### Ethics

Ethical permission was sought for the original study (18) where the collection of hospital isolates was approved by Lothian NRS Bioresource SR1094.

## Acknowledgments

This work was supported by a BBSRC Institute Strategic Programme Grant, Prevention & Control of Infectious Diseases, awarded to The Roslin Institute, a Canine Welfare Grant from the Dogs Trust (‘Advanced phage therapy for MDR *E. coli* associated with canine UTIs’) awarded to DLG, a PhD studentship supported by the School of Physics and Astronomy, University of Edinburgh awarded to IVS, and a PhD studentship supported by BBSRC EASTBIO Doctoral Training Partnership awarded to TH.

## Supplementary Figures

**Supp Fig S1: Flowchart showing the machine learning pipeline used for training the predictive models. (A)** An interaction score is calculated from the growth curves of the liquid assays. **(B)** Bacterial and **(C)** phage pangenome analyses. In each case, after the genomes are sequenced and annotated, an algorithm is used to identify all accessory genes in a binary presence/absence matrix. **(D)** Single Phage model data. Each model is built from the bacterial pangenome matrix and the interaction scores of one phage. **(E)** PanPhage model data. A single model is built by combining the bacterial pangenome matrix and interaction scores involving all bacteria and all phages. Each row contains the presence/absence of genes for one phage-bacterium pair and their interaction score. **(F)** Model training. The data is split into train and test subsets and the models are trained with the train subset. This involves repeated validation processes where the train subset is further split to reduce overfitting. The trained model is asked to predict the unseen interactions (test subset) and the predicted scores are compared to the interaction scores calculated from the growth curves (observed). **(G, H)** Performance plots comparing predicted scores (horizontal axis) and observed scores (vertical axis). Each point corresponds to one interaction in the test subset. A perfect prediction would lie on the diagonal line.

**Refer to Supplementary Figure pdf**

**Supp. Fig. S2**: Phylogenetic tree of 301 *E. coli* strains used in the study. The outer ring shows the *E. coli* phylotype (Clermont(43)) of each strain and the inner coloured ring shows the host source of each strain. The inner text ring corresponds to the bacterial strain names.

**Refer to Supplementary Figure pdf**

**Supp. Fig. S3**: Phylogenetic tree of the 31 phage used in the study (y axis) and magnified box alongside interaction data with 301 bacterial strains (x axis) (Supp Fig. S2). The heatmap shows interaction scores from growth assays. High interaction scores (greens) represent limited phage activity. Low interaction scores (reds to blue) represent high phage activity.

**Refer to Supplementary Figure pdf**

**Supp. Fig. S4**: Performance metrics of models predicting individual phage against the harmonic mean of feature distances of the predicted phage with the rest of the set. In each subfigure, data from the single-phage models (blue dots, blue dashed line), PanPhage model (orange triangles, orange dash-dotted line), and the difference between them (green squares, green dotted line) are plotted. The difference corresponds to PanPhage minus single-phage performance. The lines show linear fits of the data. The Pearson’s r and permutation p-value for each fit are (A) single-phage: r = −0.08, p = 0.7; PanPhage: r = −0.5, p = 0.01; difference: r = −0.3, p = 0.1; (B) single-phage: r = −0.1, p = 0.5; PanPhage: r = −0.4, p = 0.02; difference: r = - 0.3, p = 0.07; (C) single-phage: r = −0.1, p = 0.5; PanPhage: r = −0.2, p = 0.3; difference: r = −0.4, p = 0.05.

**Refer to Supplementary Figure pdf**

**Supp. Fig. S5**: RMSE of single-phage models and PanPhage model predicting individual phage against the activity of the predicted phage. Activity is defined as the proportion of positive observed interactions. Lines show linear fits to the data. Pearson’s r and permutation p-value are presented in the legend.

**Refer to Supplementary Figure pdf**

## Supplementary Tables

**Supp Table S1: Interaction dataset used in the study. Separate Excel file.**

**Supp Table S2:**
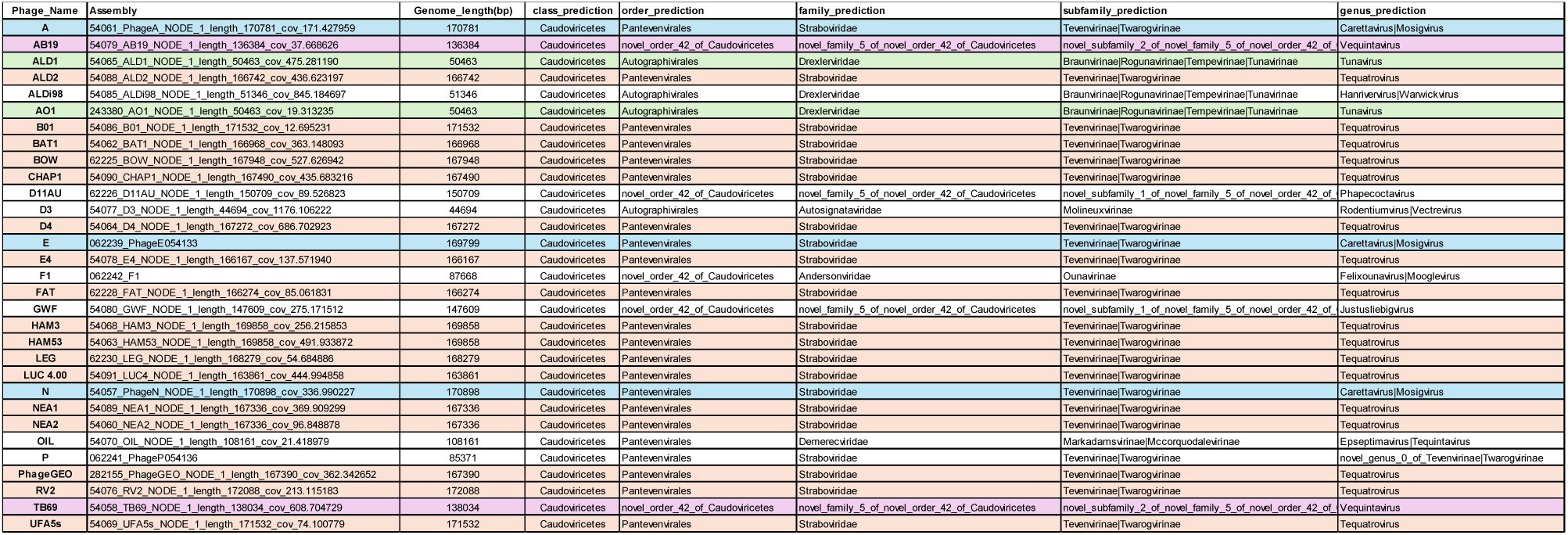
Summary statistics and predicted classifications of phage used in the study. Phage listed alphabetically. Colour coding relates to genus prediction classes. Orange – Tequatrovirus, Blue – Mosigvirus, Green – Tunavirus, Pink – Vequintavirus, White – other.

**Supp Table S3: Feature annotations. Separate Excel file**

## Supp Information (SI)1: Detailed method for machine learning models

Throughout this study, we distinguish between single-phage models and the PanPhage model. Single-phage models were trained independently for each phage and used bacterial genomic features to predict activity against individual *E. coli* isolates. In contrast, the PanPhage model was trained across all phage-bacterium interactions and incorporated genomic features from both the bacterial isolate and phage, allowing a single model to predict the interaction score for any phage-bacterium pair represented by these genomic feature sets. Wherever possible, equivalent training, feature selection and optimisation procedures were used for the two approaches, and identical train-test assignments were retained to enable direct comparison of model performance.

Models were constructed using the Gradient Boosting Regressor implementation in scikit-learn (41). The following pipeline was applied:

1. **Train-test split.** The interaction dataset was divided into training and test sets at a 3:1 ratio. Splitting was stratified by phage so that interactions assigned to the training and test sets were identical when comparing the single-phage and PanPhage approach. Because interaction outcomes were highly imbalanced for some phage, particularly those with narrow host ranges, splitting was additionally stratified by activity. Interaction scores <60 were classified as positive interactions and scores ≥60 as negative interactions (18), and the relative proportions of these classes were maintained between the training and test sets.
2. **Feature selection.** Feature selection was performed using the training data to reduce dimensionality and limit the inclusion of uninformative genomic features that could contribute to overfitting (28). The starting number of features was 16,886 bacterial genes and 1,779 phage genes. In the PanPhage model, both sets of genes were used, while in the single-phage models, only the bacterial genes were used. An initial Recursive Feature Elimination (RFE) process was performed. For this, we first trained an initial Gradient Boosting model using the complete set of features. From the fitted model, we extracted the importance of each feature, calculated as the mean decrease in variance that each feature contributes to the model, and removed the *m* least important features (*m* = 1688 for the single-phage models and *m* = 1866 for the PanPhage and LPO models). We iterated this procedure until we had 2,000 features (in the last iteration, fewer than *m* features were pruned to end with 2,000 features). Recursive Feature Elimination with a 10-fold Cross-Validation (RFECV) was then applied to the remaining features. In this process, the training data is split into 10 subsets and 10 RFE processes are carried out in parallel, each one with 9/10 of the training data (each one with different subsets). In each iteration, the performance of the fitted models is calculated by predicting on the unseen 1/10 of the training data (see “Model evaluation” and “Model performance” below). The obtained scores are averaged across the 10 models with the same number of features, and the number of features that gives the best average score is selected as the optimal number of features. Finally, a new RFE process is done, using the same step and the complete training set, to reduce the 2,000 features to the optimal number of features found. In the RFECV, we removed features in steps of 200 until a minimum of 45 was reached. Both the RFE and RFECV procedures were implemented in scikit-learn (41).
3. **Hyperparameter optimisation.** Gradient boosting hyperparameters were optimised by 25 iterations of randomised parameter search with 10-fold cross-validation using the training data. The parameter combination giving the lowest cross-validated mean squared error was selected for the final model. The parameter values considered are

a. Number of trees in the boosting ensemble: 500, 750, 1000 and 1250.
b. Maximum depth of trees: 3, 5, 10 and 20.
c. Learning rate: 0.01, 0.05, 0.1 and 0.2.
d. Subsampling fraction to be used to fit the trees: 0.8, 0.9 and 1.
e. Minimum number of samples required to split an internal node: 2, 5 and 10.
f. Minimum number of samples required at each leaf node: 1, 2 and 4.
4. **Model evaluation.** The optimised model was evaluated against the held-out test data, which was not used during feature selection or hyperparameter optimisation. Predicted interaction scores were compared with the corresponding observed scores from the wet lab experiments using the performance metrics described below.

### Model performance

To evaluate ML model performances, the root mean square error (RMSE) was calculated on the test subset predictions using the formula 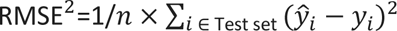, where *ŷ_i_* and *y_i_* are the predicted and observed scores, respectively, and *n* is the number of test interactions. The RMSE gives an estimation of the mean deviation between predicted and experimentally measured scores, with lower RMSE indicating greater predictive accuracy.

To additionally assess the ability of the models to discriminate between active and inactive phage-bacterium interactions, predicted and measured scores were converted to binary classifications using an interaction score threshold of 60 to define positive (score < 60) and negative (score ≥ 60) phage infections (18). Classification performance was evaluated using the F1 score and Matthews Correlation Coefficient (φ). The F1 represents the harmonic mean between the precision and recall, where precision = True Positives / (True Positives + False Positives) and recall = True Positives / (True Positives + False Negatives). The F1 score ranges from 0 to 1, with closer to 1 being better, and F1 = 1 means predicting every interaction classification correctly. The φ coefficient incorporates all four outcomes of the confusion matrix (True Positives, True Negatives, False Positives, and False Negatives) and ranges from −1 to 1, where 1 represents perfect classification, 0 performance no better than chance and −1 complete disagreement between predicted and observed classifications. The φ coefficient is defined as the Pearson correlation coefficient between the binary classification of predictions and observations, *i.e.*, 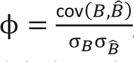, where *B̂* and *B* are, respectively, the predicted and measured infection labels in the test set, while cov and σ are the covariance and standard deviation, respectively.

To assess whether the models had true predictive capacity beyond chance, a null model was used as a baseline reference. For the RMSE measures, the null model predicts every interaction score with the average score in the training set and RMSE was calculated between these predictions and the observed test-set scores. For the F1 and φ metrics, this approach does not work because if every prediction has the same score, then all would have the same binary label (in our case, no infection). To solve this, we build a null model that predicts the binary labels drawn from a Bernoulli distribution with probability *p*, equal to the proportion of observed positive interactions in the training set. Using this approach, the φ coefficient for the null model will tend to 0 for a large number of predictions, independent of the value of *p*.

### Correlation between performance and feature-distance

We hypothesised that PanPhage predictions for an individual phage would improve when genetically related phages were represented in the training dataset. We therefore examined the relationship between model performance and the genomic distance of each phage from the remainder of the phage collection.

Pairwise phage genomic distances were calculated from the binary gene presence/absence matrix generated by pangenome analysis using the Jaccard distance. More formally, if the rows in the pangenome matrix corresponding to phage *X* and *Y* are 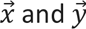, respectively, the feature distance *d*(*X*, *Y*) between them is

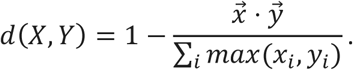

To characterise the distance between one phage and the rest of the set, we calculated the harmonic mean of the feature distances of that phage with the rest.

We studied the linear correlation of the harmonic mean of feature-distances and the performance metrics (φ, F1 and RMSE) of both the single-models and the PanPhage model, calculating the Pearson correlation coefficient *r*. To study the statistical significance of these correlations, we calculated the corresponding *p*-values by permutation testing (42).

### Feature selection statistics

The train/test split is the main source of randomness in the ML pipeline. Because the Gradient Boosting model is sensitive to which samples land in the training set, different splits will lead to different learned models. In addition, the Gradient Boosting algorithm introduces randomness when selecting between tied features when fitting the nodes. Thus, the value of the random seed will affect the genes selected through the feature selection process.

To identify genomic features consistently selected by the modelling procedure, the train-test splitting and the feature selection steps described above were repeated 128 times using different random seeds. The features retained in each iteration were recorded and ranked according to their selection frequency, calculated as the proportion of the 128 iterations in which each feature was selected.

### Leave-phage-out models

To evaluate model generalisability to phage for which no interaction data were included during training, leave-phage-out models were constructed using a leave-one-group-out approach. In each iteration, all interactions involving one phage were excluded from the training dataset and used exclusively as the test set.

Predictive performance for each excluded phage was assessed using RMSE, F1 and φ as described above. To determine whether performance on an unseen phage depended on its genomic relatedness to phage represented during training, the correlation between performance metrics and the genomic distance, between the excluded phage and the training phage collection, was evaluated using the approach described above.

## References

1. Manges AR, Geum HM, Guo A, Edens TJ, Fibke CD, Pitout JDD. 2019. Global Extraintestinal Pathogenic Escherichia coli (ExPEC) Lineages. Clin Microbiol Rev 32:10.1128/cmr.00135-18.

2. Kaper JB, Nataro JP, Mobley HLT. 2004. Pathogenic Escherichia coli. Nat Rev Microbiol 2:123–140.

3. Li X, Fan H, Zi H, Hu H, Li B, Huang J, Luo P, Zeng X. 2022. Global and Regional Burden of Bacterial Antimicrobial Resistance in Urinary Tract Infections in 2019. J Clin Med 11:2817.

4. Subramanian A. 2024. Emerging roles of bacteriophage-based therapeutics in combating antibiotic resistance. Front Microbiol 15.

5. Ibrahim R, Aranjani JM. 2026. Bacterial defense mechanisms against bacteriophages: an evolutionary arms race. Arch Microbiol 208:229.

6. Bertozzi Silva J, Storms Z, Sauvageau D. 2016. Host receptors for bacteriophage adsorption. FEMS Microbiol Lett 363:fnw002.

7. Rakhuba DV, Kolomiets EI, Dey ES, Novik GI. 2010. Bacteriophage Receptors, Mechanisms of Phage Adsorption and Penetration into Host Cell. Pol J Microbiol 59:145–155.

8. Vander Zee A, Natarajan A, Maxwell KL. 2026. Timing is everything: regulation of bacterial defences and phage counter-defences. Curr Opin Microbiol 91:102750.

9. Pourcel C, Midoux C, Vergnaud G, Latino L. 2020. The Basis for Natural Multiresistance to Phage in Pseudomonas aeruginosa. Antibiotics 9:339.

10. Aldawood E, Roberts IS. 2022. Regulation of Escherichia coli Group 2 Capsule Gene Expression: A Mini Review and Update. Front Microbiol 13.

11. Boeckaerts D, Stock M, Criel B, Gerstmans H, De Baets B, Briers Y. 2021. Predicting bacteriophage hosts based on sequences of annotated receptor-binding proteins. Sci Rep 11:1467.

12. Dunne M, Prokhorov NS, Loessner MJ, Leiman PG. 2021. Reprogramming bacteriophage host range: design principles and strategies for engineering receptor binding proteins. Curr Opin Biotechnol 68:272–281.

13. Dowah ASA, Clokie MRJ. 2018. Review of the nature, diversity and structure of bacteriophage receptor binding proteins that target Gram-positive bacteria. Biophys Rev 10:535–542.

14. Villarroel J, Kleinheinz KA, Jurtz VI, Zschach H, Lund O, Nielsen M, Larsen MV. 2016. HostPhinder: A Phage Host Prediction Tool. Viruses 8:116.

15. Gaborieau B, Vaysset H, Tesson F, Charachon I, Dib N, Bernier J, Dequidt T, Georjon H, Clermont O, Hersen P, Debarbieux L, Ricard J-D, Denamur E, Bernheim A. 2024. Prediction of strain level phage–host interactions across the Escherichia genus using only genomic information. Nat Microbiol 9:2847–2861.

16. Wei T, Lu C, Du H, Yang Q, Qi X, Liu Y, Zhang Y, Chen C, Li Y, Tang Y, Zhang W-H, Tao X, Jiang N. 2024. DeepPBI-KG: a deep learning method for the prediction of phage-bacteria interactions based on key genes. Brief Bioinform 25:bbae484.

17. Camejo PY, Rojas F, Ossa A, Hurtado R, Tichy D, Pieringer C, Pino M, Mora-Uribe P, Ulloa S, Norambuena R, Tobar-Calfucoy E, Aguilera M, Rojas-Martínez V, Cifuentes O, Sabag A, Cifuentes N, San Martín D, Infante C, Cifuentes P, Pieringer H, León LE. 2025. A machine learning approach to predict strain-specific phage-host interactions. Sci Rep 15:38249.

18. Keith M, Park de la Torriente A, Chalka A, Vallejo-Trujillo A, McAteer SP, Paterson GK, Low AS, Gally DL. 2024. Predictive phage therapy for Escherichia coli urinary tract infections: Cocktail selection for therapy based on machine learning models. Proc Natl Acad Sci 121:e2313574121.

19. Mutalik VK, Adler BA, Rishi HS, Piya D, Zhong C, Koskella B, Kutter EM, Calendar R, Novichkov PS, Price MN, Deutschbauer AM, Arkin AP. 2020. High-throughput mapping of the phage resistance landscape in E. coli. PLOS Biol 18:e3000877.

20. Miravet-Verde S, Cacace E, Mores CR, Rutschmann C, Lin C, Ruscheweyh H-J, Cuénod A, Barazzone EC, Marrec E, Vershynina K, Schumann R, Bower DJ, Schubert M, Egli A, Fiebig T, Slack E, Sunagawa S, Keys TG. 2026. In silico typing maps the natural diversity of Escherichia coli transporter-dependent capsules. Nat Microbiol 11:1217–1232.

21. Owen SV, Wenner N, Dulberger CL, Rodwell EV, Bowers-Barnard A, Quinones-Olvera N, Rigden DJ, Rubin EJ, Garner EC, Baym M, Hinton JCD. 2021. Prophages encode phage-defense systems with cognate self-immunity. Cell Host Microbe 29:1620–1633.e8.

22. Gao L, Altae-Tran H, Böhning F, Makarova KS, Segel M, Schmid-Burgk JL, Koob J, Wolf YI, Koonin EV, Zhang F. 2020. Diverse enzymatic activities mediate antiviral immunity in prokaryotes. Science 369:1077–1084.

23. Phage homing endonuclease amplifies anti-defense genes to evade bacterial immunity | Nature Communications. https://www.nature.com/articles/s41467-026-71036-4. Retrieved 17 September 2026.

24. Chihara K, Azam AH, Egorov AA, Terenin I, Hashino M, Watashi K, Horiba K, Hauryliuk V, Kiga K. 2026. Phage homing endonuclease amplifies anti-defense genes to evade bacterial immunity. Nat Commun 17:3468.

25. James T, Williamson B, Tino P, Wheeler N. 2025. Whole-genome phenotype prediction with machine learning: open problems in bacterial genomics. Bioinformatics 41:btaf206.

26. Chalka A, Dallman TJ, Vohra P, Stevens MP, Gally DL. 2023. The advantage of intergenic regions as genomic features for machine-learning-based host attribution of Salmonella Typhimurium from the USA. Microb Genomics 9:001116.

27. Lupolova N, Dallman TJ, Holden NJ, Gally DL. 2017. Patchy promiscuity: machine learning applied to predict the host specificity of Salmonella enterica and Escherichia coli. Microb Genomics 3:e000135.

28. Malajczuk CJ, Vaitekenas A, Iszatt JJ, Stick SM, Kicic A, Karpievitch YV, on behalf of PhageWA. 2026. Towards accurate artificial intelligence models for strain-level phage–host prediction. Brief Bioinform 27:bbag085.

29. King SH, Driscoll CL, Li DB, Guo D, Merchant AT, Brixi G, Wilkinson ME, Hie BL. 2026. Generative design of bacteriophages with genome language models. Science 393:eaec2657.

30. Virtanen P, Gommers R, Oliphant TE, Haberland M, Reddy T, Cournapeau D, Burovski E, Peterson P, Weckesser W, Bright J, van der Walt SJ, Brett M, Wilson J, Millman KJ, Mayorov N, Nelson ARJ, Jones E, Kern R, Larson E, Carey CJ, Polat İ, Feng Y, Moore EW, VanderPlas J, Laxalde D, Perktold J, Cimrman R, Henriksen I, Quintero EA, Harris CR, Archibald AM, Ribeiro AH, Pedregosa F, van Mulbregt P. 2020. SciPy 1.0: fundamental algorithms for scientific computing in Python. Nat Methods 17:261–272.

31. Seemann T. 2014. Prokka: rapid prokaryotic genome annotation. Bioinformatics 30:2068–2069.

32. Tonkin-Hill G, MacAlasdair N, Ruis C, Weimann A, Horesh G, Lees JA, Gladstone RA, Lo S, Beaudoin C, Floto RA, Frost SDW, Corander J, Bentley SD, Parkhill J. 2020. Producing polished prokaryotic pangenomes with the Panaroo pipeline. Genome Biol 21:180.

33. Bolger AM, Lohse M, Usadel B. 2014. Trimmomatic: a flexible trimmer for Illumina sequence data. Bioinformatics 30:2114–2120.

34. Bankevich A, Nurk S, Antipov D, Gurevich AA, Dvorkin M, Kulikov AS, Lesin VM, Nikolenko SI, Pham S, Prjibelski AD, Pyshkin AV, Sirotkin AV, Vyahhi N, Tesler G, Alekseyev MA, Pevzner PA. 2012. SPAdes: A New Genome Assembly Algorithm and Its Applications to Single-Cell Sequencing. J Comput Biol 19:455–477.

35. Nayfach S, Camargo AP, Schulz F, Eloe-Fadrosh E, Roux S, Kyrpides NC. 2021. CheckV assesses the quality and completeness of metagenome-assembled viral genomes. Nat Biotechnol 39:578–585.

36. Bouras G, Grigson SR, Papudeshi B, Mallawaarachchi V, Roach MJ. 2024. Dnaapler: A tool to reorient circular microbial genomes. J Open Source Softw 9:5968.

37. Larralde M. 2022. Pyrodigal: Python bindings and interface to Prodigal, an efficient method for gene prediction in prokaryotes. J Open Source Softw 7:4296.

38. Bouras G, Nepal R, Houtak G, Psaltis AJ, Wormald P-J, Vreugde S. 2023. Pharokka: a fast scalable bacteriophage annotation tool. Bioinformatics 39:btac776.

39. Bouras G, Grigson SR, Mirdita M, Heinzinger M, Papudeshi B, Mallawaarachchi V, Green R, Kim RS, Mihalia V, Psaltis AJ, Wormald P-J, Vreugde S, Steinegger M, Edwards RA. 2026. Protein structure-informed bacteriophage genome annotation with Phold. Nucleic Acids Res 54:gkaf1448.

40. Bolduc B, Zablocki O, Turner D, Bin Jang H, Guo J, Adriaenssens EM, Dutilh BE, Sullivan MB. 2025. Machine learning enables scalable and systematic hierarchical virus taxonomy. Nat Biotechnol 1–10.

41. Pedregosa F, Varoquaux G, Gramfort A, Michel V, Thirion B, Grisel O, Blondel M, Prettenhofer P, Weiss R, Dubourg V, Vanderplas J, Passos A, Cournapeau D, Brucher M, Perrot M, Duchesnay É. 2011. Scikit-learn: Machine Learning in Python. J Mach Learn Res 12:2825–2830.

42. Ernst MD. 2004. Permutation Methods: A Basis for Exact Inference. Stat Sci 19:676–685.

43. Clermont O, Bonacorsi S, Bingen E. 2000. Rapid and Simple Determination of the *Escherichia coli* Phylogenetic Group. Appl Environ Microbiol 66:4555–4558.

44. Hastie T, Tibshirani R, Friedman J. 2009. The Elements of Statistical Learning. Springer, New York, NY. http://link.springer.com/10.1007/978-0-387-84858-7. Retrieved 25 August 2026.

